# Cambrian origin of the arachnid brain reveals early divergence of Chelicerata

**DOI:** 10.1101/2024.02.27.582391

**Authors:** Nicholas J. Strausfeld, Frank Hirth

## Abstract

Fossils from the lower Cambrian provide crucial insights into the diversification of arthropod lineages: Mandibulata, exemplified by centipedes, insects, and crustaceans; and Chelicerata, represented by sea spiders, horseshoe crabs, and arachnids, the last including spiders, scorpions, and ticks^1^. Two mid-Cambrian genera claimed as stem chelicerates are *Mollisonia* and *Sanctacaris,* defined by a carapaced prosoma equipped with clustered limbs, followed by a segmented trunk opisthosoma equipped with appendages for swimming and respiration^2–4^. Up to the present, the phyletic status of Mollisoniidae and Sanctacariidae has been that of a basal chelicerate^2^, stemward of Leanchoiliidae, whose neuromorphology resembles that of extant Merostomata (horseshoe crabs)^5^. Here, we identify preserved traces of neuronal tissues in *Mollisonia symmetrica* that crucially depart from a merostome organization. Instead, a radiating organization of metameric neuropils occupying most of its prosoma is situated behind a pair of oval unsegmented neuropils directly connected to paired chelicerae extending from the front of the prosoma. The disposition and connections of these neuropils is the neuroanatomical signature that denotes a complete reversal of the three genetically distinct domains defining arthropod brains^6^. Thus, in *Mollisonia* the deutocerebrum is the most rostral domain with the proso-and protocerebral domains folded backwards such that tracts from the principal eyes extend caudally to reach their prosocerebral destination. These defining neuromorphological features illuminate a marine origin of Arachnida from which evolved the planet’s most successful arthropodan predators.

## INTRODUCTION

Interpreting rare neuroanatomical traces observed in Cambrian Burgess Shale-Type (BST) fossils essentially follows similar principles as those applied to interpreting neuroanatomical arrangements of extant taxa. This is because similar constraints pertain. Despite being generally compressed, fossil neuropils are volumetrically comparable to those of small euarthropods. Fossils that have split into two unequal pieces offer information about the relative location in depth of fossilized soft tissue traces just as slices of a histological specimen embedded obliquely to one side indicates vertical organization. Despite being most sharply defined at the fossil’s surfaces, neural traces are not solely resolved there: although a fossil looks opaque it may be translucent enough to allow deeper imaging beneath its surface. Images of deeper traces may appear blurred due to micro-prismatic properties of the matrix, but these effects can be mitigated by digital sharpening or using extended depth imaging in sub-millimeter steps. This corresponds to superimposing and collapsing sequential images from adjacent histological sections. Traces of fossil neuropils that appear incomplete but align with a natural pattern such as a metameric tiling reinforce the interpretation of neural traces as segmented or, crucially, as unsegmented. Overall correspondences of fossil neural traces with the densely structured neuropils of an extant taxon may provide directives for interpreting the stem affiliation of the fossil.

Here we apply those principles to examine a historical specimen (MCZ 1811) originally named *Houghtonites gracilis*^7^ *s*ubsequently assigned to the family Mollisoniida as *M. symmetrica* (Fig. 1a, b)^8^. Until now, this specimen has been claimed as an ancestral chelicerate preceding the evolution of merostome-like neuroanatomical traits defining short bodied ‘great appendage’ euarthropods such as *Leanchoilia* and *Alalcomeneaus*^2,5,9^. The overall body of *Mollisonia* essentially aligns with that of *Sanctacaris uncata*^4^ whose prosoma is equipped with at least six pairs of appendages suitable for benthic locomotion and predation as are those described from another Cambrian mollisoniid *M. plenovenatrix*^3^. Although the apparent absence of chelicerae initially suggested *Sanctacaris* as the most stemward member of Chelicerata, both *M. symmetrica* and *M*. *plenovenatrix* reveal small but robust paired chelicerae extending from the front of their prosoma^2,3^. Tagmatization into prosoma-and opisthosoma-like divisions is also demonstrated by anatomically similar members of the family Sanctacarididae^10–12^. Both sanctacaridiids and mollisoniids are morphologically distinct from short-bodied megacheirans such as *Alalacomeneaus* and *Leanchoilia* whose homonomous trunks showed no evidence of tagmatal differentiation and yet whose preserved neural traces suggest close affinity to *Limulus*^5,9,13^.

**Figure 1.**
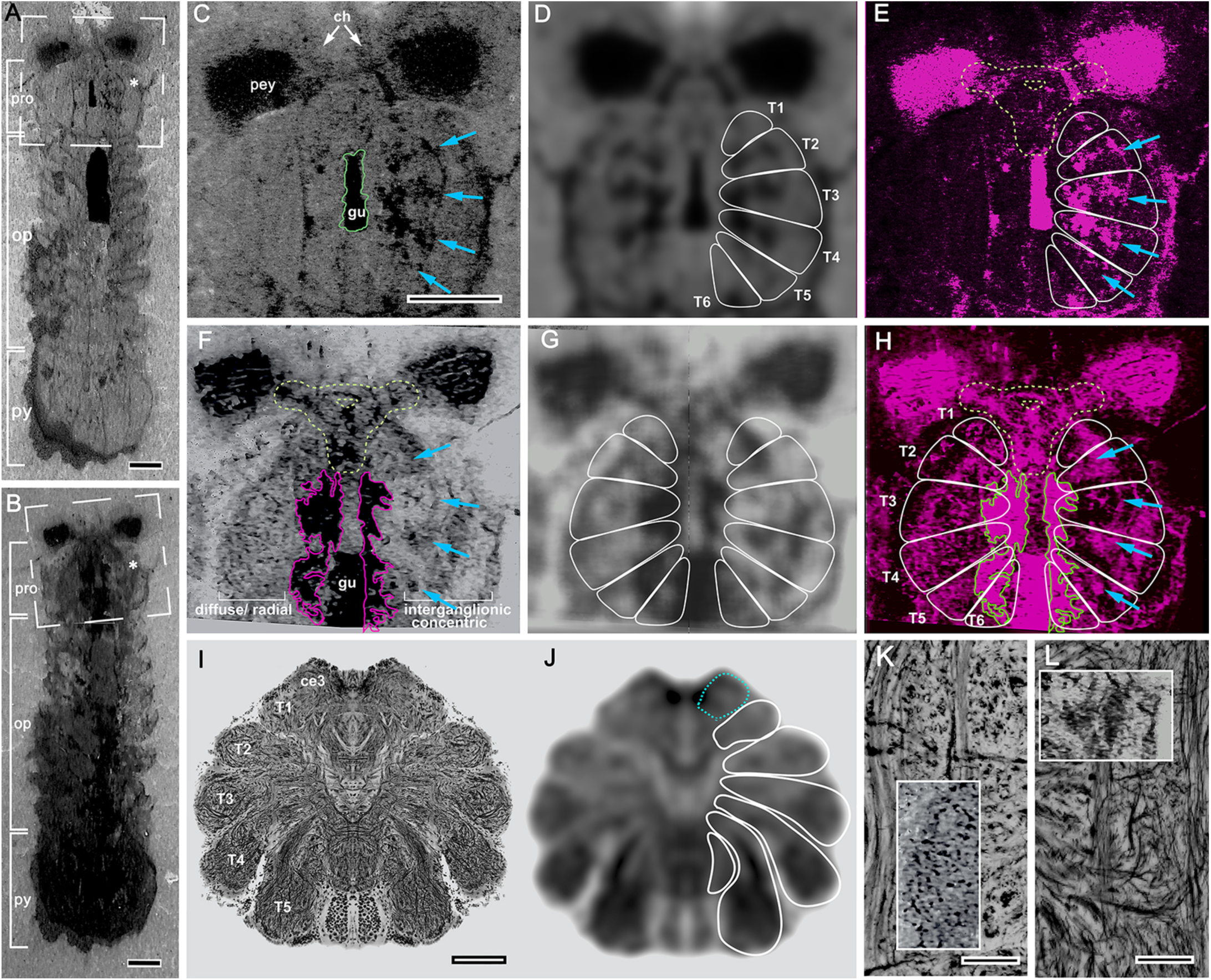
Mollisoniid prosomal nervous system conforms to the arachnid ground pattern. **(A, B)** Part (A) and counterpart (B) of specimen MCZ 1811 indicate their arachnid-like tagmatal body plan of prosoma (pro) and opisthosoma (op), terminating in a pygidium (py). Asterisks signal corresponding sides. Frames indicate orientation of the prosoma in panels C-H. Gut diverticuli are outlined in magenta or green (panels F, H). Gut with contents outlined green in panel C. Triangular areas (dashed green in F, E, H) indicate areas outside neuromeric tiling. **(C-H)** Prosoma of part (C-E) and counterpart (F-H). Both reveal profiles interpreted as neural which correspond to profiles detected as carbon deposits by energy-dispersed X-ray spectroscopy (Suppl. Fig. S4). Blue arrows in C and F indicate corresponding neural tracts and linked centers. **(D)** Mirrored Gaussian blur of the right half of panel C resolves the prosoma tiled by a fan-shape array of six neuromeres (T1-T6). **(E)** Traces of linked neuropil centers in panel C demonstrate congruence with neuromeres (panel E) as do neural traces of the counterpart (F, H). **(F)** Left-right asymmetry of neuropil traces indicate fossil tilted around the antero-posterior axis. **(G)** Gaussian blurring of the counterpart shows neuromeric congruence even of diffuse particulate processes (left half). **(H)** Highly structured neural traces in right half of panel H illustrates further congruence with neuromere tiling. **(I, J)** Reduced silver-stained prosomal neurons and their axial neuropils of the orb spider *Neoscona oaxacensis*. Gaussian blur isolates neuromeres; also shown immediately anterior to T1 (pedipalp) is the rostral position of the deutocerebrum (ce3). **(K, L)** Fossilized remnants are simpler and condensed compared with intricate networks as shown here for *Heptathela kimurai*. Features of silver-stained levels in the prosoma nevertheless align with motifs of fossil residues, as shown by particulate traces [inset width = 0.9mm in K) or layered arrangements within neuromeres [insert width = 1.25mm in L]. Scale bars; A, B = 1mm; C-H (shown in C) = 1 mm; I, J = 1mm. K, L= 50μm.

## RESULTS

We begin first by describing some general non-neural attributes of the part (MCZ 1811a, Fig. 1A) and counterpart (MCZ 1811b, Fig. 1B) of MCZ 1811 focussing on the fossilized prosoma. Its gut is situated at the fossil’s midline in MCZ 1811a. A small stretch of the anterior gut is visible in Figure 1C-E, filled by black ingested material. Absence of this material indicates soft tissue lying above the corresponding area of the counterpart, which due to its forward tilt reveals the exposed caudal continuation of the filled gut, which then continues into the opisthosoma (Fig. 1F-H). In the prosoma the gut is flanked by a system of diverticuli (outlined in Fig. 1F and 1H) identified as such by their calcium signal resolved by elemental energy dispersive spectroscopy^2^ (see acknowledgements). The most anterior length of the digestive tract within this area of the counterpart is likewise obscured by fossilized tissues preserved above it. Before describing the arrangements of fossilized neuroanatomical features two further aspects require explanation. As shown by both the part (Fig. 1C) and the counterpart (Fig. 1F), preserved traces of the nervous system are not bilaterally symmetric. This discrepancy is indicative of a bilaterian organism entombed tilted to one side, remaining so disposed throughout geological time until split into its part and counterpart along the natural grain of the embedding rock (Suppl. Fig. S1). Furthermore, even though both the part and counterpart of MCZ 1811 are taphonomically compressed, the curvature of the splitting plane allows observation of discrete components of the rostral cerebrum (Suppl. Fig. S1C, D). The counterpart’s greater opacity also indicates it is the most voluminous of the two pieces thereby accounting for most of the neural traces available for documentation here (Suppl. Fig. S2).

Both part and counterpart of *Mollisonia* (Fig. 1A, B) possess clearly defined fossilized soft tissue that we here interpret as neural. Traces near the surface of the part (Fig. 1C), when subjected to a Gaussian blur function (Fig. 1D), reveal a fan-like metameric organization of six pairs of segments against which all other neural traces can be compared and interpreted (Fig. 1C-H). However, metameric organization is absent within a triangular region of the anterior prosoma, indicated by the dashed green outline in Figs. 1E, F, H and 2A. The arrangement of segmental neuropils typifies the prosoma of extant arachnids (Fig. 1I, J), such as spiders, scorpions and sun spiders (Aranea, Scorpiones and Solifugae). Here we determine which neural traces conform to this distinctive organization of neuromeres and which traces are independent of those metameric signatures, thereby indicating the possibility of an asegmental cerebrum. Supplementary Fig. S2 demonstrates the alignment of all traceable neural residues with reference to this metameric ground pattern, which is ubiquitous to extant arachnids, here exemplified by a silver-stained nervous system of the orb spider *Neoscona oaxacensis* (Fig. 1I, J).

We first consider the part (Fig. 1A) in which a sharply defined tract on the right side connects four condensed clusters of branched aggregates (Fig. 1C). This organization corresponds to the trajectory of a less distinct but clearly visible system in the counterpart (Fig. 1F). In both panels these correspondences are indicated by the four blue arrows (Fig. 1C, E for the part, and Fig. 1F, H for the counterpart]). To what extent do other neural traces conform to this metameric ground pattern? Considering the counterpart, demonstrating a metameric distribution of its radially oriented particulate traces might be seen as challenging (Fig. 1F). Nevertheless, the general outwardly directed orientation of these small, elongated profiles approximately align with the fan-like disposition of the prosoma’s neuromeres. That the observed profiles indeed reflect the neuroarchitecture of segmental ganglia is demonstrated by organization in the T1 appendicular neuromere. Its two distinct levels of differentiation (Fig. 1H) is further shown in Fig. 2A where the medial part of the T1 neuromere, indicated by the blue arrow, corresponds to the base of the T1 ganglion in the corresponding neuromere of the extant spider *H. kimurai* (Fig. 2B, corresponding arrow to that of T1 in panel A). The distal arrangements in the mollisoniid T1, albeit simpler and condensed, correspond to the more elaborate processes and neuropils in the outer part of the corresponding aranean T1 (Fig. 2B). Although the basal ganglionic components of neuromeres T2 and T3 are resolvable (blue arrows) more caudal ones can only be inferred (open blue arrows) with particulate traces in the left side of the counterpart representing neuromeres T4-T6. These arrangements contrast with more structured features in the right side of both the part and counterpart (Fig. 1C, E; F, H). Neural traces reflecting discrete segmental aggregations, as in Fig. 1E and Suppl. Fig. S2 A-C, are distinct from traces that extend across neuromeres that provide wide-field networks as shown in the right side of Fig. 1H and in Suppl. Fig. S2D-F. These distinctions correspond to the functional organization of circuits in living arachnids. Shown by intracellular injection of dyes in araneans, discrete arrangements of motor neuron arborizations and receptor neuron terminals relating to individual ganglia, as also indicated in fossil traces (Suppl. Fig. S2C), are located more ventrally and medially in the prosomal nervous system than are connections that coordinate information across ensembles of neuromeres^14,15^. The latter are mediated by widely distributing interganglionic arrangements^16^ such as indicated by fossil traces depicted in Suppl. Fig. S2F. Overall, even though fossilized remnants are never as intricate as in histological preparations due to the obvious clumping of some fossil traces, they align with neural architectures revealed in silver-staining sections of living arachnids (Fig. 1K, L).

**Figure 2.**
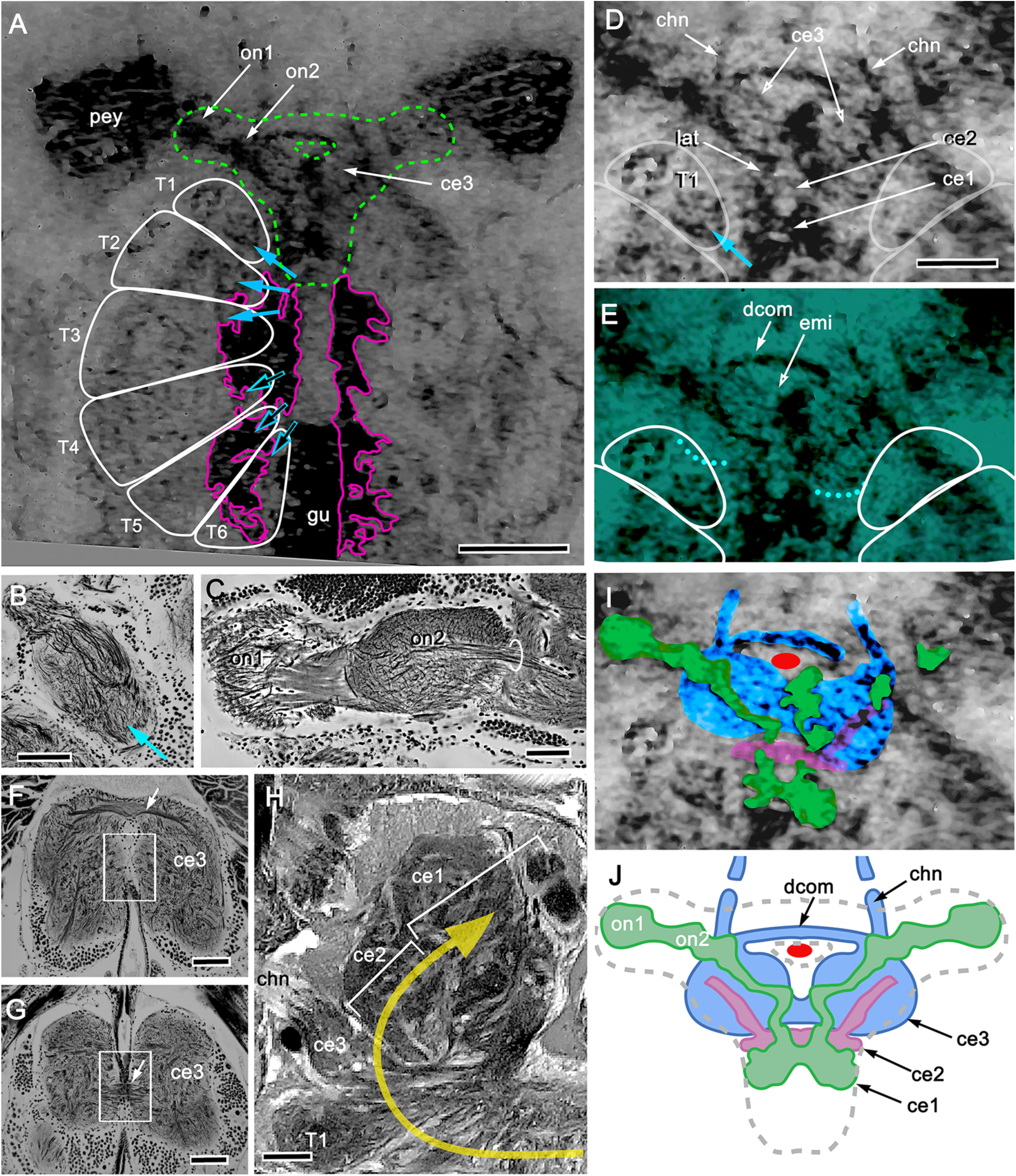
Cerebral arrangements of *M. symmetrica* are outside metameric tiling. **(A)** Counterpart of MCZ 1811. Magenta outlines gut diverticuli disposed alongside the visible gut (gu) and extending partially across where more anteriorly the gut lies beneath neural tissue. The ganglion-like roots of neuromeres T1–T3 (solid blue arrows) are well defined, those in T4–T6 are obscured (open arrows) beneath diverticuli. **(B)** The T1 neuromere of *H. kimura* is organized into proximal motor and distal sensory neuropil levels. These correspond to two levels in the mollisoniid T1, enlarged in panel D). **(C)** Peripheral visual centers typifying extant arachnids (here *Delena cancerides*) send their central relays (circled) to distant central neuropils. This arrangement corresponds to the first and second visual neuropils (on1, on2 in A) that extend from the edge of the mollisoniid principal eye (pey) and its medially directed visual nerve. **(D)** Enlargement of the area outlined in green in A reveals the nerve leading from the principal eye’s visual neuropils and transversing one of a pair of oval centers. These are here identified as ce3 due to their forward extending nerve into the chelicerae (cheliceral nerves; chn). Two circumscribed regions lie immediately behind ce3: a small transverse bridge (ce2) followed by a larger domain (ce1) in which terminates the principal eye’s lateral optic nerve (lat). **(E)** A digital green filter offers clear resolution of the ce3 domains (posterior margins indicated by dotted lines) and their linking dorsal commissure (dcom), which indicates the location of the endomesodermal interface (emi). **(F, G)** Two levels of the ce3 domain of extant *H. kimura* demonstrate the upper (F) and lower (G) commissures typifying the euarthropod deutocerebrum^23^. Boxed areas indicate passage of the gut. The emi would be located immediately anterior to the lower commissure. **(H)** Lateral view of the cerebral domains in a histological section of the orb spider *Tetragnatha* sp. The arrow indicates the antero-posterior neuraxis in which the ce1 (prosocerebrum) is the bona fide anterior-most domain, whereas, as a result of the brain bending back over itself, the ce3 (deutocerebrum) has become the anterior-most cerebral domain according to the body axis. **(I)** Color codes attributed to the reversed cerebral domains ce3, ce2, and ce1. **(J)** Interpretive reconstruction of the mollisoniid cerebrum. Scale bars: A = 0.5mm; B, C = 100 μm; D (also for E, I) = 250 μm; F, G, H =100μm.

The one part of the prosomal nervous system that cannot be reconciled with overall metameric organization is the approximately triangular area situated in front of the T1 neuromeres (Fig. 1E, F, H; 2A (dashed outline). Within this area several well-defined tracts and swellings are associated with the principal eyes. Enlargement of this area (Fig. 2A) reveals two connected optic neuropils that are isolated from the cerebrum proper and are thus distinct from an integrated optic lobe, such as reported for the mandibulate *Fuxianhuia protensa* and typifying mandibulates in general^17, 18^. The organization shown in Fig. 2A typifies the arachnid visual pathway where, as in spiders (Fig. 2C), the first integrative optic neuropils are distant from their higher central visual centers, which are embedded in the proso-and protocerebra^19, 20^.

But where in *Mollisonia* are these components of the cerebrum? To identify them requires a brief visit to a living arachnid. As shown in Fig. 1I, the aranean prosoma comprises five distinct neuromeres T1-T5 on each side. In addition, there is a pair of neuropils immediately adjacent and rostral to segment T1. These neuropils are designated ce3 to denote the arachnid brain’s deutocerebral domain. In merostomes, pycnogonids and mandibulates, this domain is posterior to the protocerebrum, as it is in Leanchoiliidae^5,6,18,22,23^. In the generic arachnid represented by Fig. 1I, segment T1, which serves the pedipalps and is homologous to the mandibular tritocerebrum, is immediately posterior to ce3. However, in Fig. 1I there is the glaring absence of the ce2 and ce1 cerebral domains; namely, the protocerebrum and prosocerebrum. Scrutiny of *Mollisonia* reveals the same peculiar arrangement (2A, D, E). Its T1 domain is likewise the homologue of the pedipalp ganglion with T2-T6 relating to 5 additional pairs of appendages (the mollisoniid T6 appendage is absent in crown araneans, there being only four pairs of walking legs). As in extant Aranea, the anterior flank of the mollisoniid T1 abuts the border of an immediately adjacent oval domain, which, with its heterolateral partner, is extreme rostral situated outside the metameric area of the prosomal nervous system (Fig. 2D, E). Enlargement of these domains, labeled ce3 in Fig. 2D, shows the one on the left overlapped by the second optic neuropil and its tract extending medially to its central destination in ce3. These oval domains are not part of the visual system, however: they give rise to forwards-extending nerves denoted as ‘chn.’ that reach into the chelicerae. This organization – paired rostral lobes connected to the chelicerae – is the crucial anatomical signature informing that these anteriormost domains must be the deutocerebrum that has adopted an extreme rostral cerebral location. Even though the fossil has undergone considerable flattening, the rostral position of these neuropils corresponds to the unique arachnid disposition of the deutocerebrum as shown half a billion years later by the extant example in Figs. 1I, 2H. As we emphasize above, in all euarthropods except Arachnida the deutocerebrum is located behind the first two domains of the cerebrum (the proso-and protocerebra, thus ce1 and ce2) as demonstrated by the innervation of the “great appendages” of the stem merostome Leanchoiliidea^9^ and by developmental studies of *Limulus* and pycnogonids^21,22^. Only in arachnids is the ce3 domain thrust forwards to lead the ce2-and ce1 domains, which are folded backwards obscuring the most anterior part of the gut (Fig. 2H).

A cardinal feature of euarthropods is that the two lobes of the deutocerebrum are connected by a ventral and a dorsal commissure,^23^ and it is these that signify the location of the endoderm-mesoderm interface (emi) denoting the transition of the foregut to the hindgut. This occurs immediately anterior to the junction of the deutocerebrum and the T1 segment^6^. Here, with the uniquely rostral location of the deutocerebrum (ce3), this interface occurs posterior to the dorsal commissures linking the two lobes of ce3 as indicated in Fig. 2E. This accords with the organization of the ce3 domain of *H. kimurai*, where the gustatory passage penetrates through the deutocerebrum between the dorsal and ventral commissures as shown in Fig. 2F, G. The reversed arrangement of cerebral domains in an arachnid is best demonstrated seen from the side, as in Figure 2H of the orb spider *Tetragnatha* sp., which illustrates the stacked cerebral domains in their reverse order -ce3, ce2, ce1-corresponding to the three cerebral domains ce3, c2 and ce1 in the fossil *Mollisonia* (Fig. 2D) whose rostro-caudal sequence is discernible despite compression of the fossil (see Suppl. Fig. S1). As shown in Fig. 2D-E, the optic neuropil tract passes caudally to reach the third cerebral domain (ce1), lying across the midline above the gut, which is hidden beneath it. As has been already emphasized, the connections between peripherally located optic neuropils and their distant integrative centers in ce1 (and ce2) is a defining trait of the arachnid brain^19,20^. In extant arachnids a second optic nerve from secondary eyes terminates in the intermediate heterolateral domain ce2^19^. Whether this also occurs in *Mollisonia* is still unresolved. The mollisoniid *M*. *plenovenatrix* clearly has two pairs of large eyes^3^, whereas *M. symmetrica* is reported to have just one pair^2^. Yet observation of one side of the surface of MCZ 1811a suggests that there may be a simple eye immediately adjacent to the larger facetted eye (Suppl. Fig. 3). Its presence is further suggested by a faint profile indicating that ce2 is supplied by a short afferent pathway (Fig. 2D, I). Each domain, color-highlighted for clarity in Fig. 2I, provides the interpretative depiction shown in Figure 2J. This organization is further compared in Fig. 3 with that of an extant aranean *Phiddipus appachiensis,* aligning the corresponding endomesodermal interface of both arachnids with that of the “linear brain” of the leanchoiliid nervous system (Fig. 3B, C).

**Figure 3.**
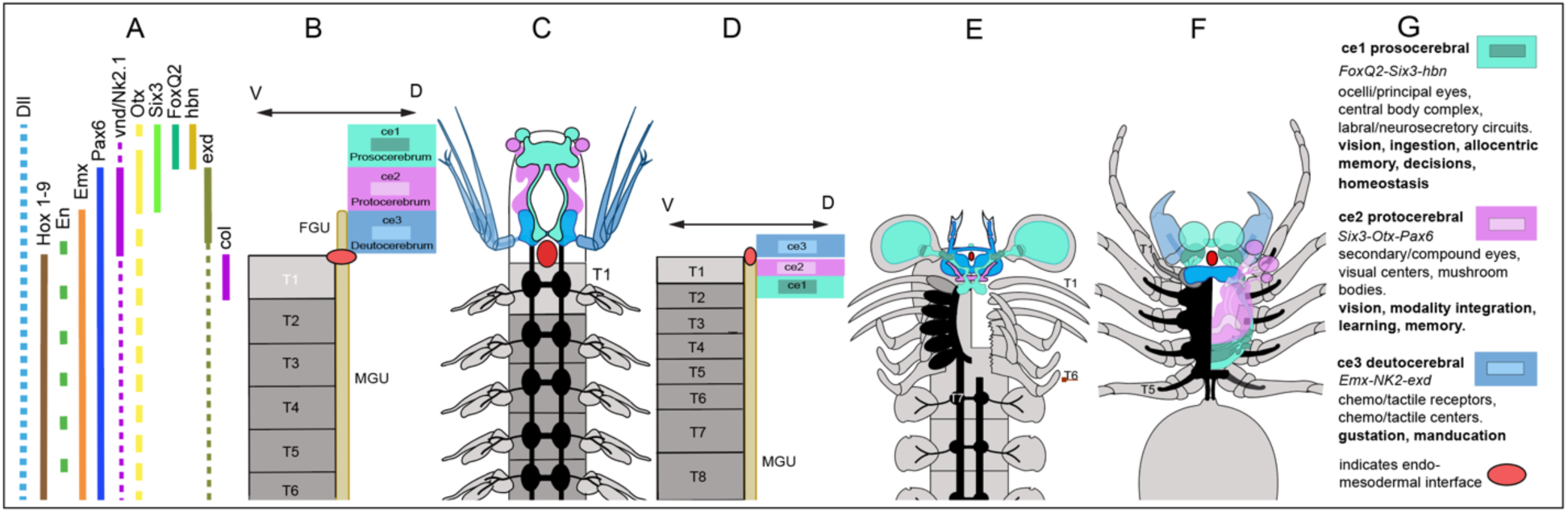
The linear cerebrum of *Alalcomenaeus* and the reflected cerebrum of Arachnida. (**A**) Combinatorial expression of homologous genes across extant taxa (for source references see *9*) defines the three domains of the asegmental cerebrum (ce1 prosocerebrum; ce2 protocerebrum; ce3 deutocerebrum) with first trunk segment T1 defined by anterior Hox gene expression and *collier (46)*. The junction between the asegmental cerebrum and the segmental trunk coincides with the endomesodermal interface (red ovoid). (**B, C**) Linear brains exemplified here by the megacheiran *Alalcomenaeus*, aligned along the anterior-posterior axis with the cerebrum situated dorsally above the foregut (FGU) and subsequent trunk ganglia beneath the midgut (MGU), as schematically shown with its deutocerebral nerves supplying the chelate appendages that define the most caudal domain ce3. (**D**) Schematic of the reflected arachnid brain and segmental ganglia (gray, T1-T8 etc.). (**E**) Top-down view into the prosoma and first three opisthosomal segments (T7 and following) of *Mollisonia*, with the carapace and right half of the neuromeres removed to expose the underlying prosomal appendages (T1-T6) schematized here with reference to M. *plenovenatrix* ^3^. The reflected cerebrum, typified by *Mollisonia* and defining extant Arachnida, locates the deutocerebrum (ce3 domain) forward as the most rostral domain (according to body axis) from where it supplies nerves to the paired chelicerae. (**F**) The reflected cerebrum is essentially unchanged in extant arachnids, here represented by the brain of a salticid aranean in which the ancestral prosomal appendages are reduced to paired pedipalps and four pairs of walking legs. (**G)** Genetic code of brain domains, the resulting domain-specific neural centers and associated sensory traits that apply across extant Euarthropoda and Onychophora^6^.

## DISCUSSION

Elucidating the ancestral organization of the arthropod head and its brain has been called “the endless pursuit”^24^, in part due to the striking diversification of cephalic morphologies, but also because of the assumption that partitions of the underlying cerebrum reflect the organization of the encapsulating subdivided exoskeleton. Yet early studies on nervous system embryogenesis inform that the embryonic mandibulate cephalon is not segmental despite its tripartite organization^25^. Recent studies have confirmed this both for arachnids and mandibulates^26,27^. Understanding the identity and evolution of these asegmental domains is therefore crucial in defining cephalic traits that distinguish mandibulates from chelicerates and, within the latter, arachnids from merostomes and pygnogonids. In considering this it is useful to bear in mind that across Euarthropoda the cerebrum is genetically defined as three unique neural domains (ce1-ce3), each specified by the combinatorial activity of highly conserved homeobox genes^28–30^ (see also Fig. 3G). In crown euarthropods, each cerebral domain is characterized by its unique genetic code as are the neuropil constituents of each domain and its associated sensory attributes (Fig. 3A, B, G). These neural arrangements of extant euarthropods align with identifiable fossilized soft tissues interpreted as neural in the cerebrum of *Alalcomenaeus* (Fig. 3C) and *Mollisonia* (Fig. 3E). However, the crucial distinction between them is that in *Alalcomenaeus* (and living merostomes and pycnogonids) the cerebrum is linearly arranged along the antero-posterior body axis (ce1 to ce3), (Figure 3B, C), whereas in *Mollisonia* and living arachnids the cerebrum is bent backwards, with the deutocerebrum leading the reversed proto-and prosocerebrum order (ce3 to ce1) along the body axis (Fig. 3D-F).

If the same unique combination of conserved gene expression patterns defines the integrative centers that distinguish each of the three cerebral domains, irrespective of whether they are linear or reflected^30,31^ this begs the question of what advantage does the reflected brain confer on arachnids. This has not been debated despite the elaborate arrangement in arachnids of segmental neuromeres that govern sensory perception and motor control having direct access to computations mediated by neural circuits of the ce1 and ce2 domains. The first domain, ce1, is defined by its elaborate arcuate body connected to the principal eyes; the second domain, ce2, is defined by its chemo-and motion detection mushroom bodies that integrate sensory information from the secondary eyes and the chelicerae^19,20^. Both domains provide the most immediate connection to the metameric neuropils of the prosoma whose elaborate and complicated interconnections have no parallel in either merostomes or mandibulates^14–16^. It is reasonable to infer that these unique arrangements in mollisoniids have been retained throughout half a billion years of arachnid dominance as the planet’s most successful arthropodan predators.

That the reflected cerebrum is a defining trait of Arachnida brings into focus the significance of short-bodied Megacheira represented by *Alalcomenaeus* and *Leanchoilia* that are ancestral to Merostomata^5,9^. These stem chelicerates are characterized by their linear cerebra as are Mandibulata,^18^ a distinction that answers the question of whether *Mollisonia* should have the status of an inclusive upper stem ancestor of total Chelicerata as has been proposed^2^ or, as shown here, the status of an exclusive upper stem ancestor of total Arachnida that diverged early from the chelicerate stem, as schematized in Figure 4. It is notable that the first *Limulus*-like fossil preserved with its entire serial central nervous system demonstrates that the location of its cheliceral ‘ganglia,’ which are the paired deutocerebral lobes, is posterior to its proso-and protocerebra^32^.

**Figure 4.**
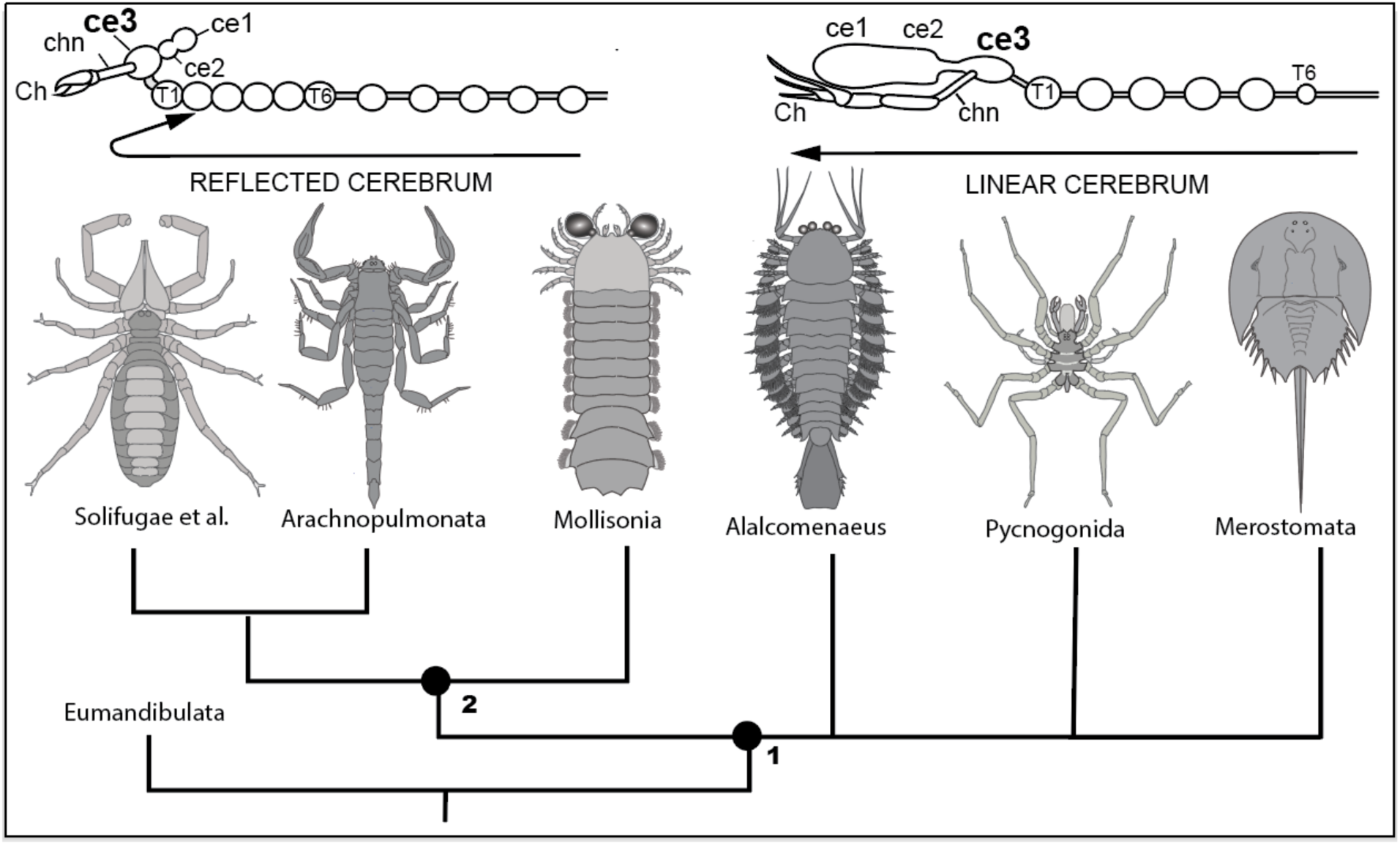
Schematic indicating major relationship within Chelicerata *sensu lato*. Two fossil total-group chelicerates, *Mollisonia* and *Alalcomenaeus,* exemplify divergent character states of the three asegmental domains of the cerebrum and their invariable order, namely, linear (ce1-ce2-ce3) or reflected (ce3-ce2-ce1) according to the antero-posterior body axis. The lineages *Alalcomenaeus*, Limulidae and Pycnogonida participate in a four-way polytomy (node 1) with the clade total-group Arachnida at node 2. These relationships resonate with recent phylogenomic analyses of all living chelicerate orders^35^. The present data identify the megacheiran *Alalcomenaeus*^2^ as the stem taxon closest to Pycnogonida and Xiphosura/Merostomata, and *Mollisonia* as the stem taxon closest to extant Arachnida, thus revealing a marine origin of Arachnida and total group Chelicerata. As part of the character state “reflected brain”, the ce3 domain lies most anterior with regards to the A-P body axis, its short cheliceral nerve (chn) reaching to the chela (Ch). In the linear brain, ce1 is the most rostral brain domain with regards to the A-P body axis, whereas ce3 is the most caudal brain domain which provides an extended cheliceral nerve reaching its deutocerebral appendages.

Here we have shown that the differences in organization of cerebral traits, in particular the relative position of the deutocerebrum, indicate that the divergence of Arachnida and Merostomata had occurred by the mid-Cambrian. Whereas the megacheiran *Alalcomenaeus*^5^ is a stem taxon ancestral to the lineage providing Xiphosura, *Mollisonia* is the stem taxon closest to extant Arachnida and indicates a marine origin of that class, *sensu* Ballesteros et al.^33^ while enlivening the question whether terrestrialization of marine arachnids occurred just once, as currently accepted^34^.

## METHODS

### Fossil provenance

The part and counter part of *Mollisonia* (MCZ 1811a and MCZ 1811b) are curated and stored at Harvard Museum of Comparative Zoology (HMCZ).

### Fossil analysis

Dr. Javier Ortega-Hernandez (HMCZ) arranged access to specimens MCZ 1811a and b for photographic documentation that provided the data discussed herein. Specimens were photographed wet with crosspolarization. Images were taken mainly at medium to low light intensity with exposure times ranging from about 0.5-3 seconds. Images were acquired using a Zeiss Axiocam 208 color camera mounted on a Zeiss Stemi 305 microscope linked to a laptop computer running Zen 3.5 software. A Zeiss CL6000 light source with flexible dual branch light guides was equipped with twistable polarization filters. Red or green gel Wratten filters were occasionally used for mixing polarized and direct illumination. Selected enlargements of high-definition files were processed using Adobe Photoshop 24.6.

### Histological microscopy

The extensive collection of silver-stained nervous systems representing major arachnid groups was generated in the N.J.S. laboratory and provided comparison with trace neuropils described herein. Images of the reduced silver-or osmium-stained cerebrum of aranean brains were taken using a JenaOptic Progress C3 digital camera fitted to a Leica Diaplan microscope equipped with plan neofluar oil immersion objectives.

## ACKNOWLEDGEMENTS

We thank Drs. Javier Ortega-Hernández for providing facilities and equipment at the Harvard Museum of Comparative Zoology and Dr. Rudy Lerosey-Aubril (HMCZ) for assistance in setting up the relevant photographic instrumentation. Dr. Ortega-Hernández kindly provided the elemental maps of MCZ 1811b^2^, showing calcium identification of gut diverticuli and carbon evidence for nerve tracts and putative neuropil (Suppl. Fig. S4). Dr. Wulfila Gronenberg (University of Arizona) kindly provided the sectioned brain of *Tetragnatha* sp, used for Figure 2H. Dr. Greg Edgecombe (Natural History Museum, London. U.K.) kindly advised us regarding the description of Figure 4. Our very special thanks go to Dr. Camilla Strausfeld for her advice and critical editing of the manuscript.

## Author contributions

N.J.S. and F.H. originated the project; N.J.S examined and photographed the fossil. F.H. ascribed published gene expression data to the described panarthropods. N.J.S. and F.H. wrote the manuscript.

## Supplementary Figures

**Supplementary Figure S1.**
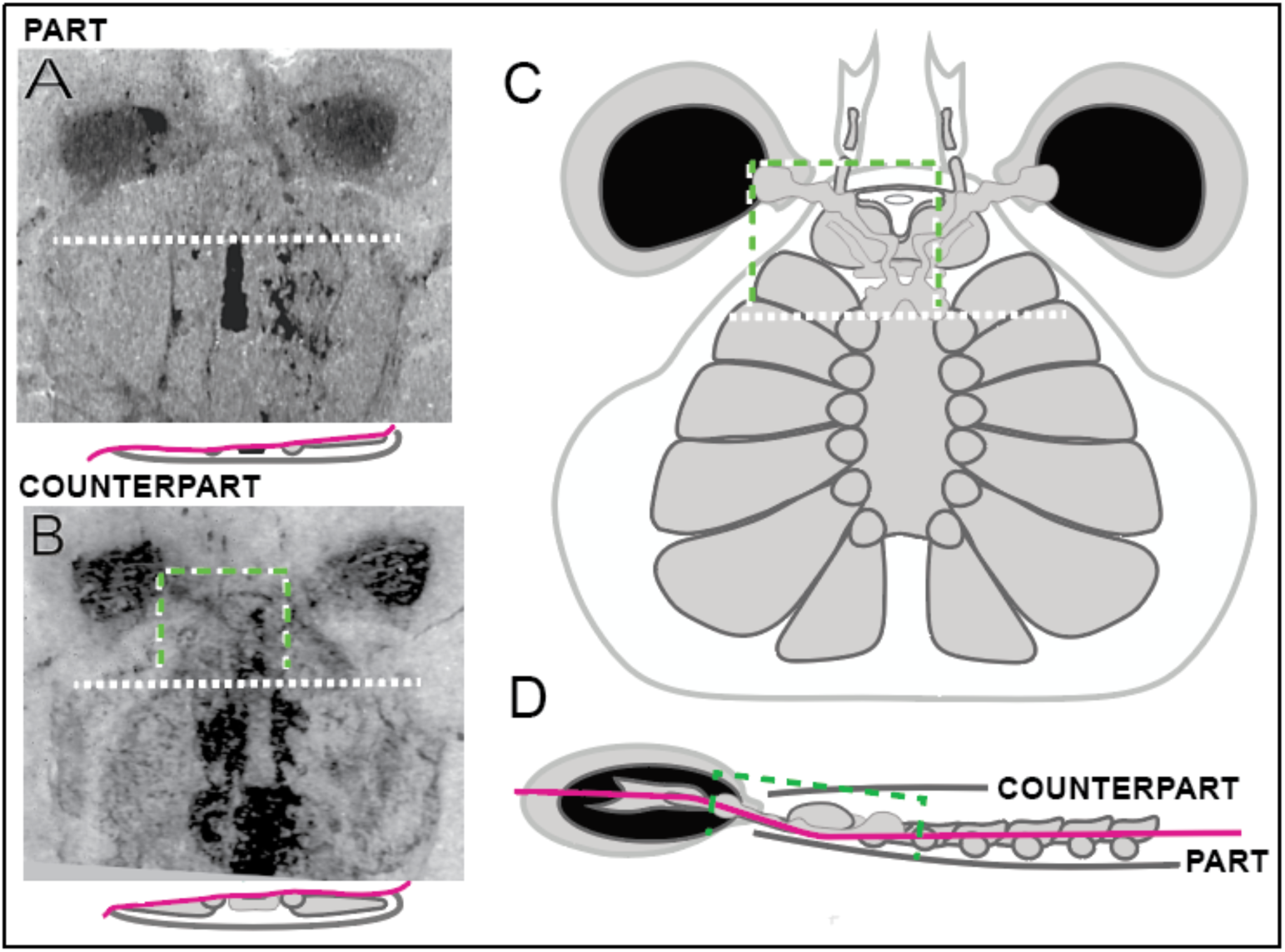
Disposition and fracture of the part and counterpart of *M. symmetrica* specimen MCZ 1811. Reconstructed brain and segmental neuromeres have relied on a fracture plane whose left-right tilt allows scrutiny of different levels of fossilized neural traces **A**, **B**, and hence reconstruction of metameric organization (**C**). The curved antero-posterior course of fracture (**D**) exposes medial rostral neuropils of the cerebrum (green dashed bracket), as described in the main body of the text.

**Supplementary Figure S2.**
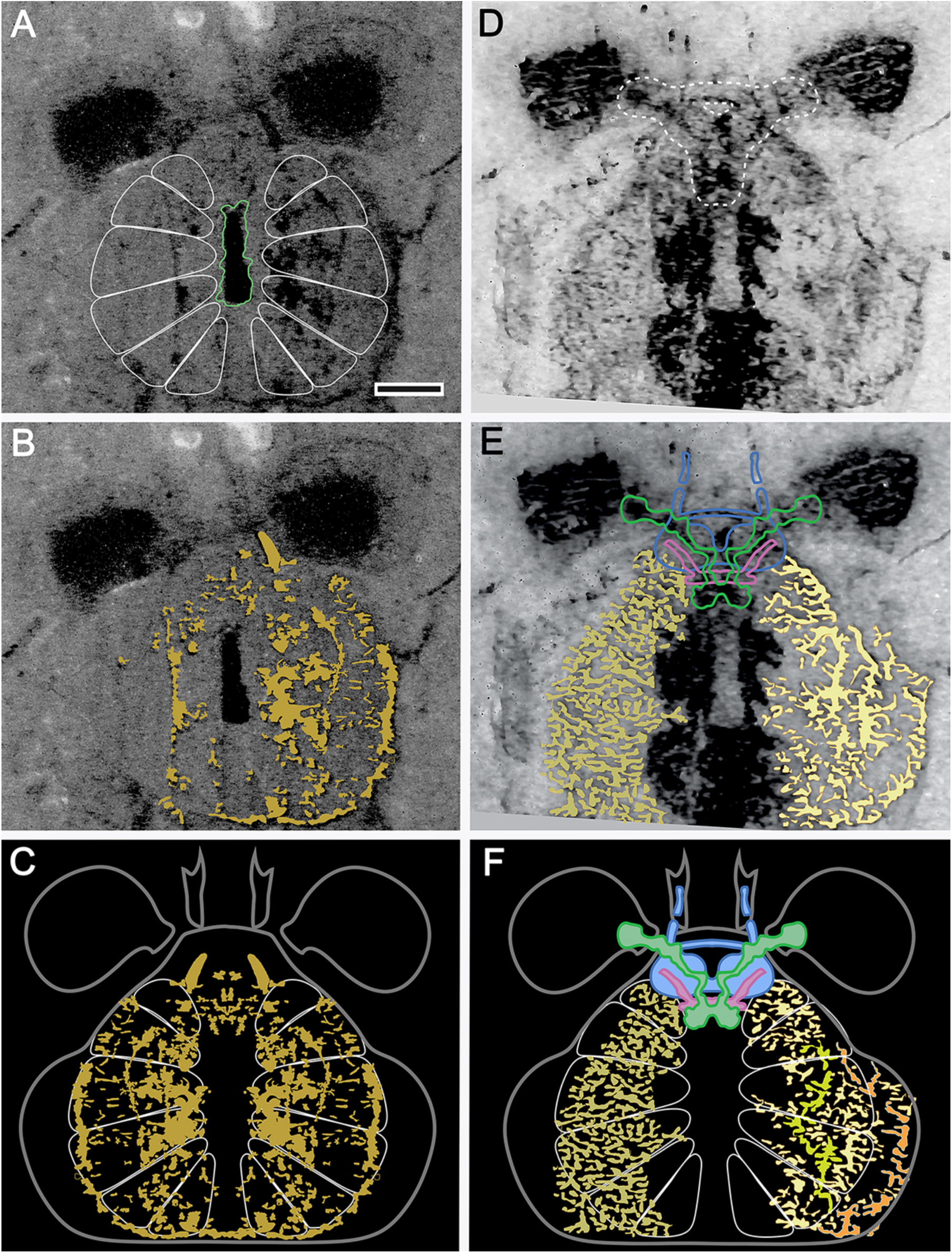
Fossilized neural traces of *M. symmetrica*. (A-C) Metameric divisions of the prosoma (B) superimposed on neural traces in the part (B) demonstrate their contribution to segmental neuromeres. The part provides the most ventral traces illustrating the concentric organization of segmental arrangements closer to the midline, whereas multisegmental traces extend through the periphery of each neuromere. (D-F) The asymmetry of counterpart provides two populations of traces: on the left many particulate traces generally fan outwards constrained to the metameric divisions; on the right two arborizations \provide clusters of processes that conform to the neuromeres. A peripheral trace does not; its disposition suggesting a more dorsal distribution. The inclusion of the reconstructed cerebrum in panel E, F indicates the relative size of the asegmental brain vis-a-vis the extensive neuropils of the prosomal segmental nervous system. Scale bar in A, also for B, D, E = 0.5mm.

**Supplementary Figure S3.**
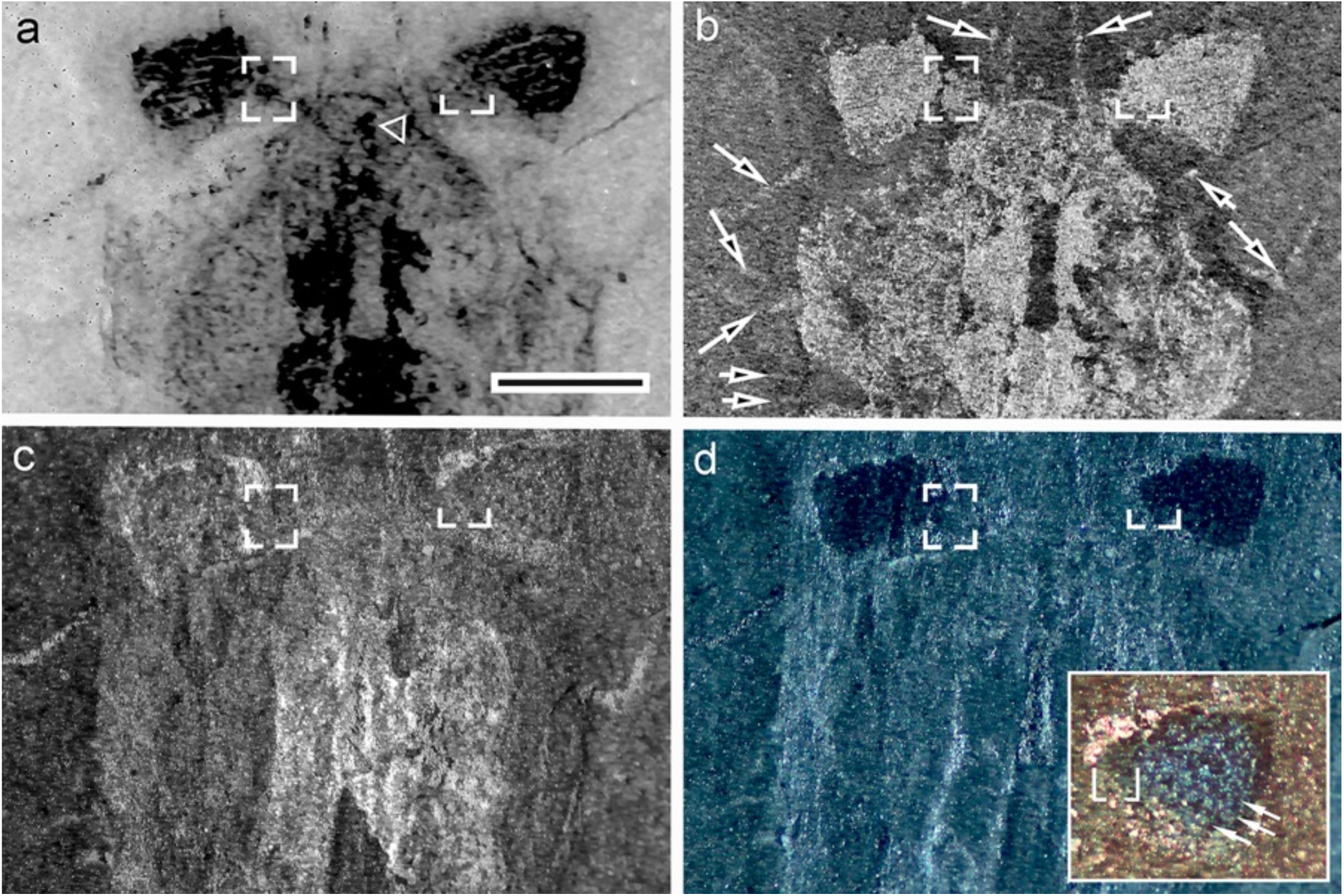
Ambiguities of eye morphologies and eye number of *M. symmetrica*. Evidence that *M. symmetrica* possesses two immediately adjacent eyes is not immediately apparent. (**A**) Neural traces in the counterpart MCZ 1811b resolve two large dark areas with some streaks as evidence for one pair of eyes associated with the outer optic neuropil (boxed in left half). This neuropil is missing from the right side (expected position bracketed) and appears to have been displaced possibly at the time of death to a medial location indicated by the open triangle. (**B**) Low-angle, cross-polarization of MCZ 1811b reveals reflective features including the outer optic neuropil (boxed). Its equivalent contralateral area also is strongly reflected despite the absence of its neuropil. Remnants of prosomal appendages (arrowed) are also illuminated. Notably, the edges of the paired chelae extend forwards from the prosoma (open arrows). (**C**) MCZ 1811a: Bright low-angle illumination of the immediate surface of the water-immersed part (C) hints at a teardrop profile for the right eye (frames and brackets correspond to those in panels A, B). **(D)** Extreme low-angle, cross-polarized illumination of surface features (water immersion) showing the large (principal) eye and medially adjacent to it a small surface structure (bracket). The inset shows an enlargement under ambient red with cross-polarization to resolve rows of facets (arrowed) in the principal eye and a small possibly receptor surface suggesting a small secondary eye (bracketed). Scale bar in A for panels A-C=1 mm.

**Supplementary Figure S4.**
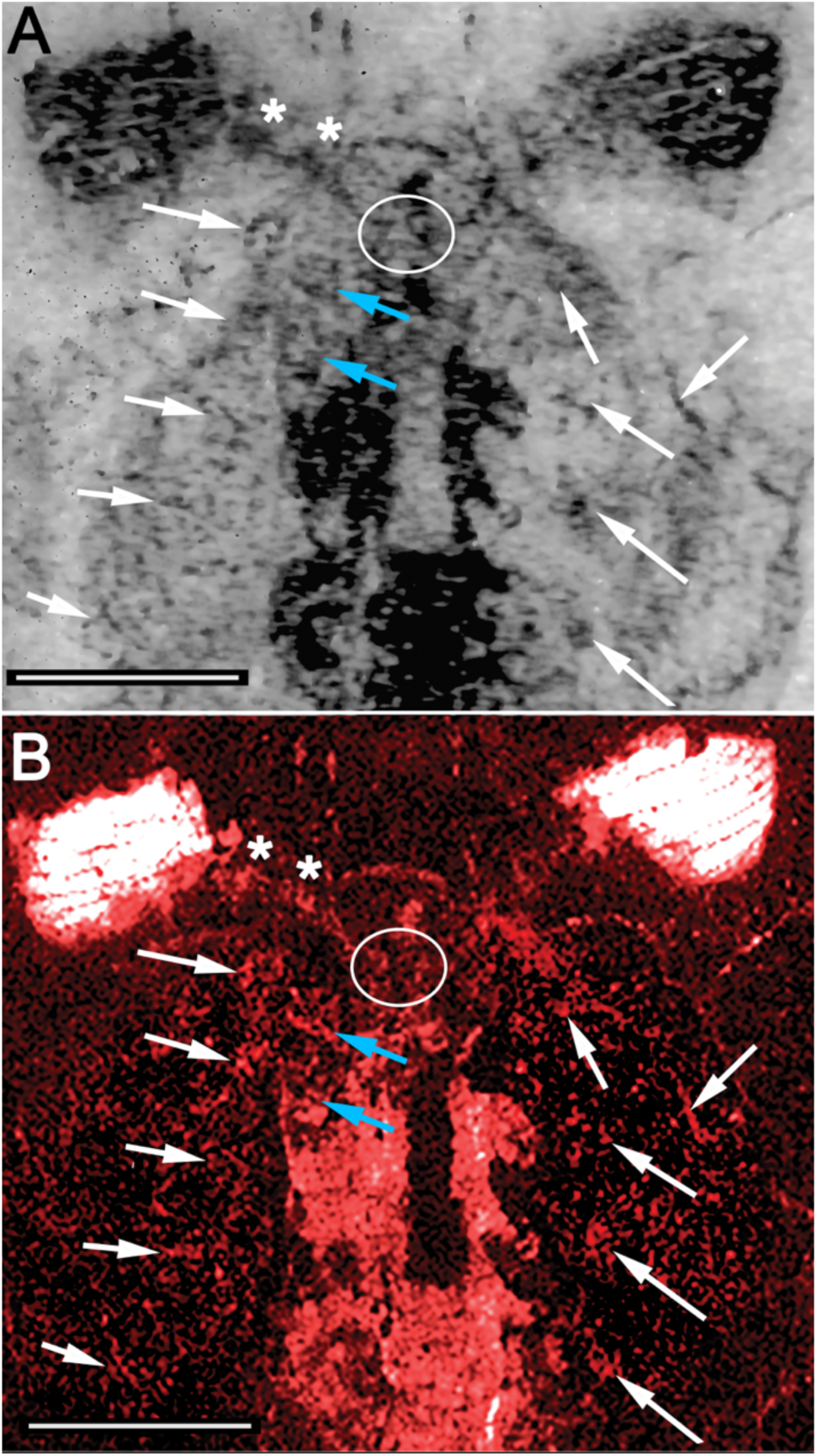
Photographic correspondences of neural traces and their carbon signatures of *M. symmetrica* (MCZ 1811b) See also ref. 2 and acknowledgments. **(A)** Neuropils and tracts in the counterpart of *M. symmetrica*, MCZ 1811b, resolved photographically. **(B)** Carbon map obtained by energy-dispersed X-ray spectroscopy (EDS). Arrows indicating a selection of corresponding features (e.g. blue arrows indicating T1 and T2 neuromeres) interpreted as fossilized nervous system. Oval in A illustrates greater clarity of the photographed image of the optic nerve (asterisks) compared with its ‘noisy’ carbon representation. Comparison of the photographic image and the EDS image also resolves false positives in EDS scan suggesting less reliability in terms of neuroanatomical analysis than photography. Scale bar= 1mm.

